# Microbial Potlatch: Cell-level adaptation of leakiness of metabolites leads to resilient symbiosis among diverse cells

**DOI:** 10.1101/2020.11.06.370924

**Authors:** Jumpei F Yamagishi, Nen Saito, Kunihiko Kaneko

**Author notes:** **For correspondence:**, (KK); (NS).

## Abstract

Microbial communities display extreme diversity, facilitated by the secretion of chemicals that can create new niches. However, it is unclear why cells often secrete even essential metabolites after evolution. By noting that cells can enhance their own growth rate by leakage of essential metabolites, we show that such leaker cells can benefit from coexistence with cells that consume the leaked chemicals in the environment. This leads to an unusual form of mutualism between “leaker” and “consumer” cells, resulting in frequency-dependent coexistence of multiple microbial species, as supported by extensive simulations. Remarkably, such symbiotic relationships generally evolve when each species adapts its leakiness to optimize its own growth rate under crowded conditions and nutrient limitations, leading to ecosystems with diverse species exchanging many metabolites with each other. In addition, such ecosystems are resilient against structural and environmental perturbations. Thus, we present a new basis for diverse, complex microbial ecosystems.

## Introduction

In microbial communities, extremely diverse species or strains coexist (***Lozupone et al., 2012**; **Curtis et al., 2002**; **Datta et al., 2016***), while secreting and exchanging hundreds of (essential) metabolites such as amino acids and sugars (***Baran et al., 2015**; **Ponomarova et al., 2017**; **Embree et al., 2015***), vitamins (***Zengler and Zaramela, 2018***), and nucleotides (***D’Souza et al., 2014**);* an archaeon transfers lipids and possibly even ATP (***Morris et al., 2013**; **Huber et al., 2012**).* Even when only a single resource is available, varieties of species coexist, and the dominance of a single fittest species that excludes all others is seldom observed (***Rosenzweig et al., 1994**; **Goldford et al., 2018**).* In contrast, according to the competitive exclusion principle proposed by Gause (***Gause, 1932***), the number of coexisting species in a common environment cannot exceed the number of available resources in the environment (***Hardin, 1960**; **MacArthur and Levins, 1964**).* Actually, the observed diversity may not be inconsistent with this principle because metabolite secretion and exchange can, in principle, create new niches and thereby allow for the coexistence of diverse microorganisms. This has been recognized both experimentally (***Goldford et al., 2018**; **Zelezniak et al., 2015***) and theoretically (***Goyal and Maslov, 2018**; **Marsland III et al., 2019***).

Nonetheless, the evolutionary origin(s) of secretion and exchange of essential metabolites that support microbial coexistence and symbiosis is still enigmatic. A constructive laboratory experiment revealed that stronger cells (i.e., cells with higher glutamine synthetase activity) coexist with weaker cells, via leakage of glutamine synthesized by the *former (**Kashiwagi et al., 2001***). Lenski et al. stressed the importance of chemical leakage by proposing the black queen hypothesis (BQH), a theory on the evolution of metabolic dependency based on gene loss (***Morris et al., 2012**; **Morris, 2015***). However, these studies did not examine whether leakage is beneficial for leaker cells. Leakage is simply assumed to be inevitable because of the permeability of their cell membranes (***Morris, 2015**; **Großkopf et al., 2016**; **Zomorrodi and Segrè, 2017***), albeit disadvantageous it may be. If this is the case, there is no reciprocity between the leaker cells and the other cells; then, why have the leaker cells not evolved to decrease the leakiness and not dominated the ecosystem?

A recent theoretical study showed that leakage of even essential metabolites can promote the growth of the leaker cells even in isolated conditions (***Yamagishi et al., 2020***). This phenomenon is termed as *leak advantage.* Indeed, microorganisms leak a variety of essential metabolites even in isolated conditions, including central metabolic intermediates (***Paczia et al., 2012***). This leak advantage can explain why leakage of even essential metabolites is preserved or acquired through evolution (***Braakman et al., 2017***), and may provide a new perspective on metabolite-mediated microbial ecology.

In the present paper, we examined whether and how cell-cell interactions mediated by secreted metabolites can lead to stable coexistence of diverse microbial species (or strains or mutants), rather than the dominance of a single fittest species. We first show that “leaker” and “consumer” species (i.e., cells that benefit by leaking some chemicals and those that benefit by consuming them, respectively) can immediately develop a mutualistic relationship. We then explore the conditions under which such “leaker-consumer mutualism” is observed and stable.

Based on the idea of leaker-consumer mutualism, we will further show that when each coexisting cell species optimizes its own growth (which may result from adaptation within a generation or evolution over generations), the coexistence of diverse species is achieved and the overall growth rate of the microbial community is enhanced. This novel scenario for symbiosis among diverse species will explain why the single “fittest” species does not dominate as a result of evolution. Furthermore, we will show that systems with exchange of metabolites among diverse species are resilient against external perturbations. Finally, we discuss the possible relevance of the present results for experimental characterization of the resilience of microbial ecosystems.

## Model

Let us consider a situation where cells that contain *n* kinds of chemical components (metabolites and enzymes) coexist in a common environment (Fig. 1A), as seen in previous studies (***Furusawa and Kaneko, 1998**; **Kaneko and Yomo, 1994**; **Yamagishi et al., 2016***). The state of cell *α* is expressed by the concentrations of the *n* components, 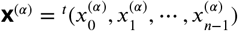. In each cell, chemical *i* is synthesized and decomposed by a set of intracellular reactions at the rate 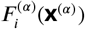 and is exchanged with the environment at the rate 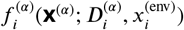, where 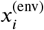 represents the concentration of chemical *i* in the environment. Here, out of *n* chemical components in each network, *N*_enzyme_ chemicals are “enzymes” which could be a catalyst or product of each reaction, while the rest of the chemicals, termed as “metabolites”, could be diffusible (if 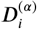 is positive) substrates or products of each reaction.

**Figure 1.**
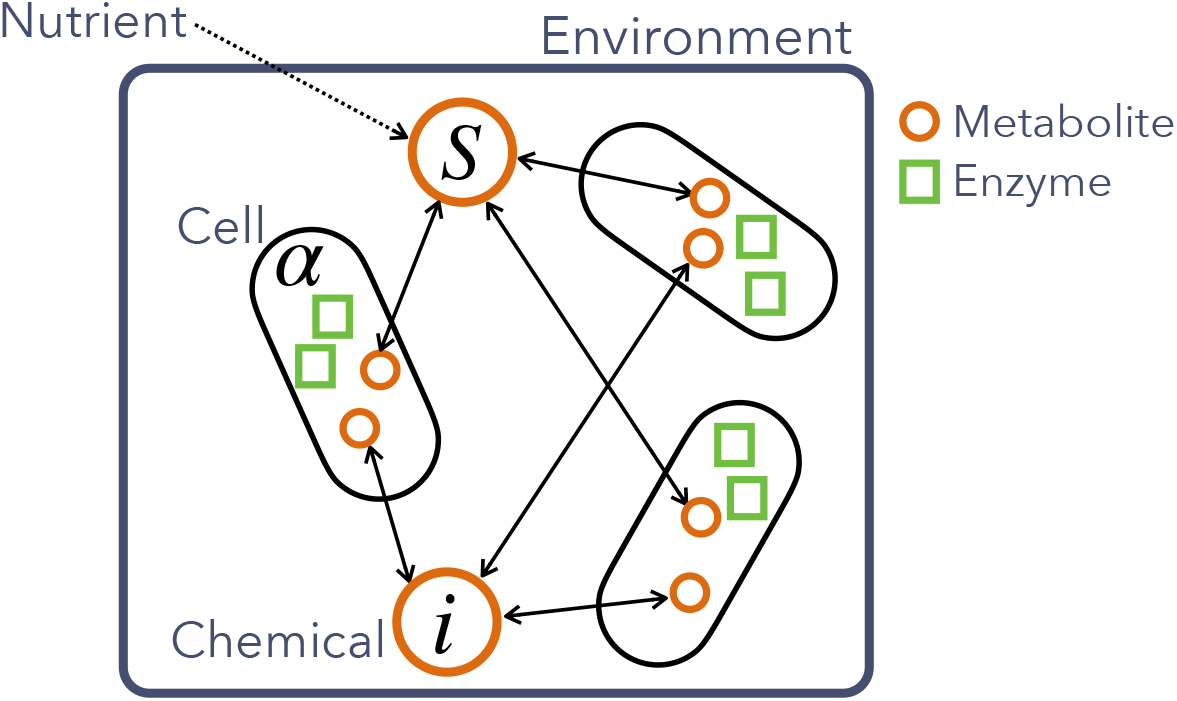
Schematic illustration of the model for microbial community with metabolite exchange. Each cell species has a chemical reaction network that transforms a single nutrient *S* transported from the environment for cell growth. The nutrient *S* is supplied to the environment from the exterior at the rate 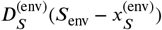. Among *n* chemicals, metabolites (orange circles) are diffusible and exchanged by coexisting species via the environment, while enzymes (green squares) are not.

The temporal evolution of the concentration of chemical *i* in cell 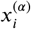, is generally written as where

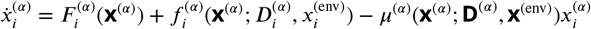

where, 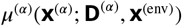 is the volume growth rate of *α*th cell, and the third term represents the dilution of each chemical owing to the increase in cellular volume. We here discuss the case of passive diffusion, where the flow rate of chemical *i* is given by 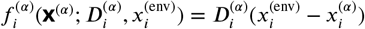. 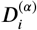 is a non-negative parameter characterizing the flow rate of each metabolite *i*, which we call the diffusion coefficient. If 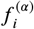 is positive, then chemical *i* flows into cell *α* from the environment, and if it is negative, *i* is leaked out. Note that we define *leak-advantage chemicals* for species *α* such that an (infinitesimal) increase in their leakage promotes the growth of species *α* (***Yamagishi et al., 2020***).

To account for cell-cell interactions due to the transport of chemicals through the environment, the time evolution of the external concentrations **x**^(env)^ is given as

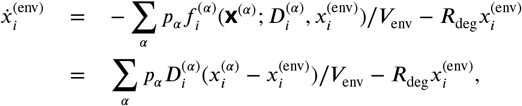

when chemical *i* is not a nutrient. If chemical *i* is a nutrient, it is supplied into the environment via simple diffusion, so that the term 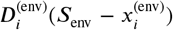 is added to the right-hand side of the above equation. In the external medium, the secreted components slowly degrade or flow out of the medium at the rate 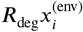. The volume of the environment relative to the total volume of all the coexisting cells is designated as *V*_env_.

In addition, the population fraction of cell species *α*, given by *p_α_*, evolves according to the equation

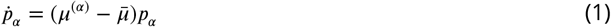

where *μ*^(*α*)^ is the growth rate of each cell species, and 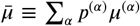 is the average growth rate of all species (***Kaneko, 2016***).

We then investigated the steady state of the population dynamics of multiple species with different reaction networks, and examined whether they can coexist in a common environment.

## Results

### A simple example of leaker-consumer mutualism: symbiosis between two species

In order to exemplify the leaker-consumer mutualism, we first consider the simplest situation: symbiosis between two cell species in which the leaker cells secrete an essential metabolite and the consumer cells consume it to facilitate their growth.

For the sake of simplicity, the network structure of the example in ***Yamagishi et al. (2020)***—a simple reaction network that consists of substrate ***S***, enzyme ***E***, ribosome rb, metabolites ***M***_1_ and ***M***_2_, biomass BM—is adopted both for the leaker and consumer cells (Fig. 2A). The equations for the reactions are given below, and the rate constants given below are different for the leaker and consumer cells:

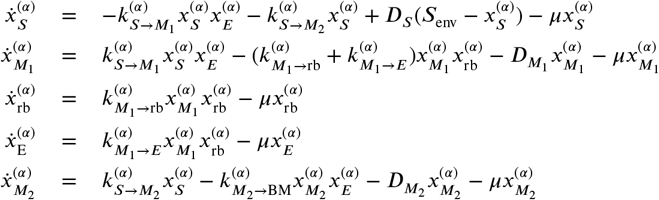

where the growth rate is defined as the rate of synthesis of biomass BM from its precursor ***M***_2_, such that 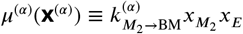. Although all chemicals—***S***, ***E***, rb, ***M***_1_, and ***M***_2_–are necessary for the growth of cells with this reaction network, certain levels of leakage of metabolite ***M***_1_ can promote the growth of cell ***α*** with a relatively large rate constant 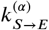. In this example of symbiosis between two cells, the rate constants of the leaker and consumer cells are setat 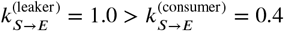; thus, the leakage (uptake) of ***M***_1_ is beneficial only for the former (latter) (Fig. 2A).

**Figure 2.**
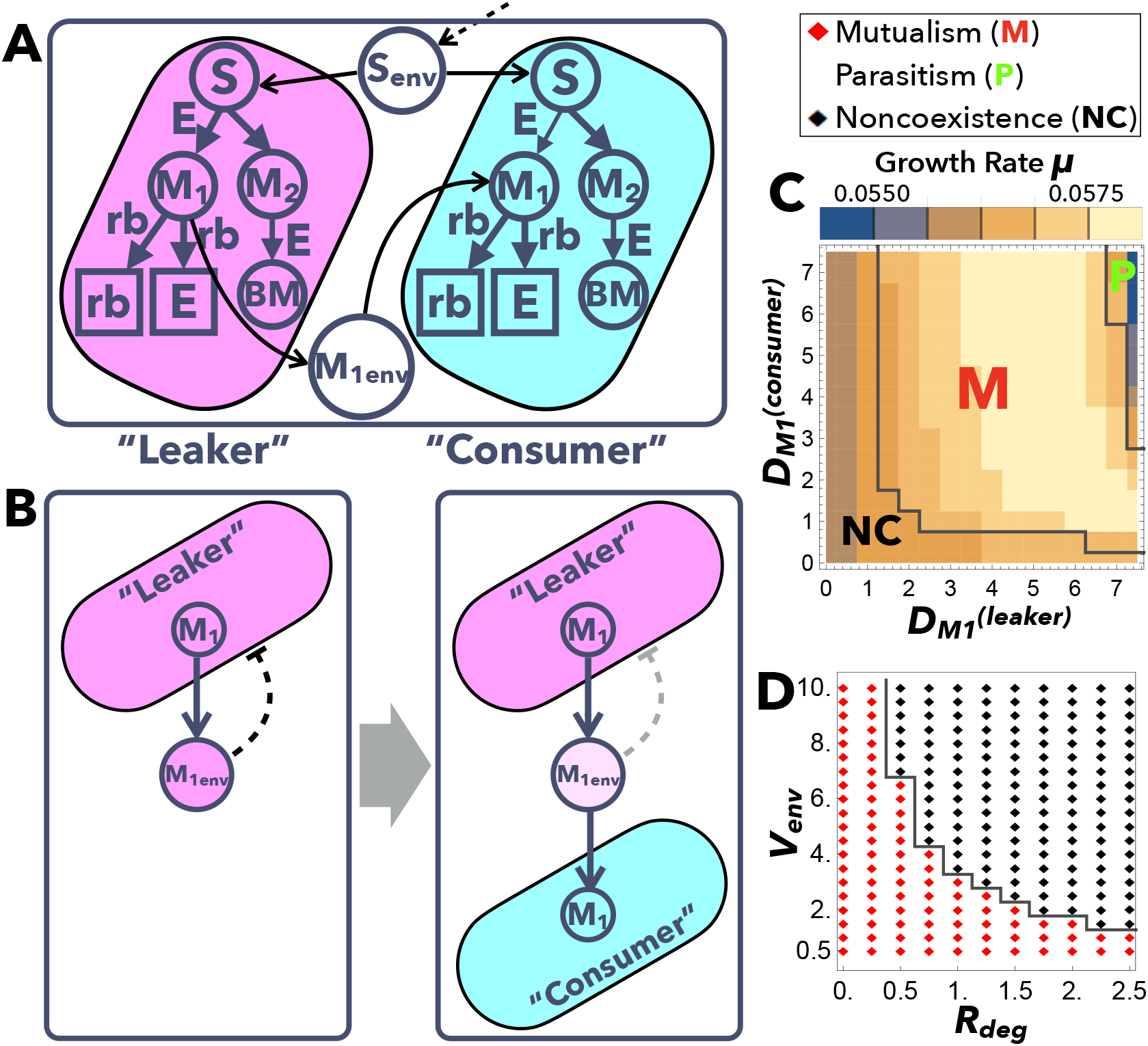
Example of leaker-consumer mutualism between two cell species. (A) Schematic illustration of the mutualism between the leaker (left) and consumer (right) cells. Both have the same network structure as shown in ***Yamagishi et al. (2020)*** with different rate constants. (B) Schematic illustration of leaker-consumer mutualism. When only a leaker cell is present, the secreted chemical accumulates in the environment and inhibits further secretion (left). The coexistence of other consumer cells that consume the leaked chemicals is beneficial for both cells because it relaxes the accumulation of the chemical in the environment (right). (C) Phase diagram of symbiosis depending on 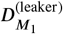 and 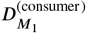. Regions M (red), P (green), and NC (black) are delineated by gray lines and represent mutualism, parasitism, and noncoexistence, respectively. The environmental parameters are set as ***V***_env_ = 1 and ***R***_deg_ = 1. The color denotes the growth rate ***μ***, where a brighter color corresponds to a higher ***μ. μ*** with 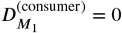 is just the growth rate of the leaker cells in isolated conditions, 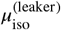; in region M, it is smaller than ***μ*** at the corresponding 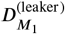 value in the panel. (D) Phase diagram of symbiosis depending on the environmental parameters, ***R***_deg_ and ***V***_env_. The diffusion coefficients of ***M***_1_ are fixed as 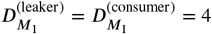. Red and black diamonds are delineated by gray lines and represent mutualism and noncoexistence, respectively. The rate constants are set as: 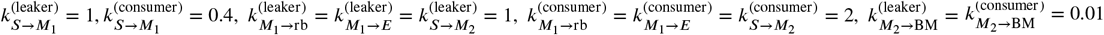, so that the leaker’s growth rate in isolation 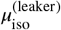 with optimal diffusion coefficient 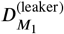 is higher than the consumer’s growth rate in isolation 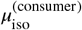 with 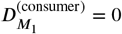. The other parameters are set as 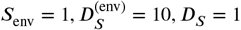.

Our numerical simulations show that the mutualism between the leaker and consumer cells is achievable under certain conditions, as schematically illustrated in Fig. 2B). When only the leaker cells that gain a leak advantage by secreting a chemical are present in the system, the secreted chemical accumulates in the environment so that further secretion becomes difficult due to the loss of the concentration gradient (see the right panel in Fig. 2B). Since the leaked chemical is an essential metabolite and not a waste product, other species can use them for their own growth in most cases. This consumption of the leaked chemical by other cell species is also beneficial for the leaker cells, as it reduces the accumulation of the leaked chemicals in the medium (see the left panel in Fig. 2B).

Hence, the leaker and consumer cells coexist through this leaker-consumer mutualism, when diffusion coefficients range within certain values indicated as Region M (Mutualism) in Fig. 2C. Under such conditions, the growth rates of the two cells are equal, which is higher than that of each cell species put in an isolation condition (see 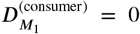 in Fig. 2C which correspond to the growth rates of the leaker cells in isolation). However, when 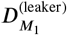 and/or 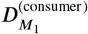 are small, the growth rate of the consumer cells cannot increase to that of the leaker cells, and thus, only the leaker cells exist in the environment (Noncoexistence; Region NC in Fig. 2C). In contrast, when 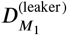 and 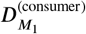 are large, the leaker and consumer cells can still coexist, but their growth rate is lower than that observed when leaker cells grow in isolation. Thus, parasitism, rather than mutualism, is realized in such cases (Region P in Fig. 2C). This is because excess leakage of necessary chemical ***M***_1_ is disadvantageous to the leaker. Figure 2C, however, also indicates that the fastest growth is achieved by mutualism between the two types of cells. Accordingly, if both cells adaptively alter their diffusion coefficients, parasitic coexistence is excluded, and mutualistic coexistence is expected to emerge. Notably, such cell/individual-level growth optimization via adaptive changes of diffusion coefficients of each cell spontaneously leads to optimal growth at the community/ecosystem level.

Additionally, Fig. 2D reveals that the leaker-consumer symbiosis is achieved if ***R***_deg_ and ***V***_env_ are not very large, that is, if the secreted chemical is efficiently transported to the other cell. However, when the degradation rate ***R***_deg_ or environment size ***V***_env_ is too large to allow for sufficient accumulation of the secreted metabolite in the environment, the cel Is no longer coexist, and only the leaker cell survives (see also ***Kashiwagi et al., 2001***).

### Symbiosis among randomly generated networks because of cell-level adaptation

To investigate the possibility of leaker-consumer mutualism and symbiosis among more cell species with diverse chemicals, we further considered a model including a variety of cell species with randomly chosen catalytic networks. The transport of chemicals from one cell species to another can be bidirectional if their membranes are permeable to diverse chemicals, which may lead to a complex symbiotic relationship. For simplicity, we mainly considered reaction networks including only catalytic reactions such as ***i*** + ***k*** → ***j*** + ***k*** with a catalyst ***k*** (the simplest multibody reactions; see also ***Yamagishi et al. (2020))*** and equal rate constants (set at unity), and only a single nutrient (chemical 0) is supplied externally.

We first generated a “species pool” containing ***N*** = 50 randomly generated networks, and these species were added into the environment one by one. At the time of the addition, the invading species optimizes the diffusion coefficients of non-nutrient metabolites to maximize its growth rate under the environmental state **x**^(env)^, where other cell species exist before the invasion. After the addition of the new species, the population dynamics of Eq. [1] are computed over a sufficiently long period ***T***, until the population distribution reaches a steady state (species with population ratio smaller than the threshold value *p*_min_ = 2.5 × 10^-4^ will be eliminated). After this procedure, each surviving species gradually and simultaneously alters its diffusion coefficients of non-nutrient metabolites over period ***T*** so that its own growth rate increases. This process of invasion and cell-level adaptation is repeated for all ***N*** species (see Supplementary Material for details).

To examine whether symbiosis among cells with randomly chosen networks can be achieved as a consequence of cell-level adaptation of leakage and uptake, the above model was numerically studied. As shown in Fig. 3A, invasions can occasionally reduce the number of cell species in the environment, but generally, the number and the growth rate of coexisting species increase. Consequently, multiple cell species could steadily coexist by exchanging multiple metabolites even under a single-nutrient condition, even though the cell species have different growth rates in isolation conditions (Fig. 3BC). Because of this adaptive metabolite exchange, the growth rates of the different species become equal, and higher than that of each species in isolated conditions (Fig. 3C). Thus, symbiosis at the community level is achieved by cell-level adaptation.

**Figure 3.**
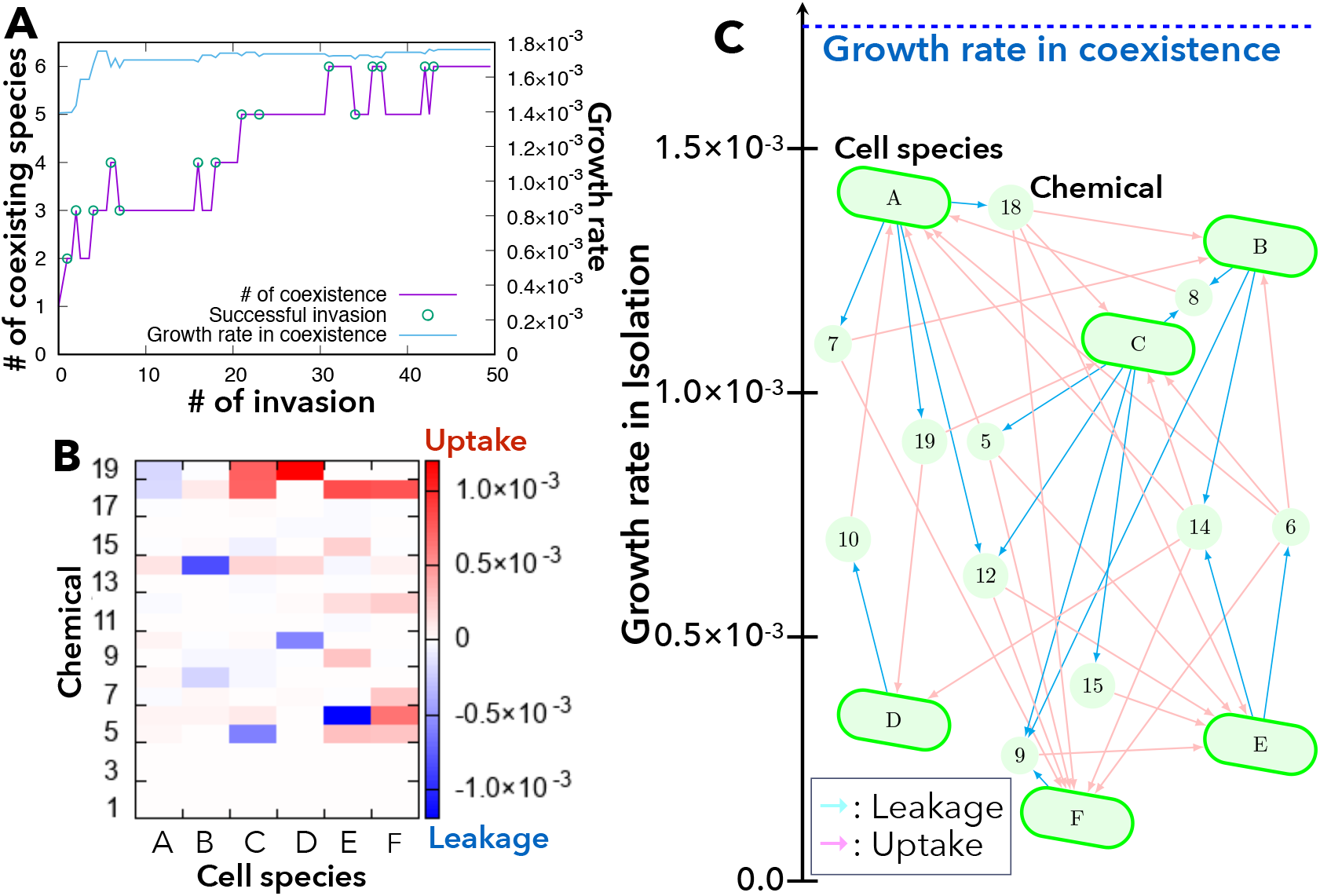
An example of symbiosis with metabolic exchange via the environment. (A) Time series of the number of coexisting species through successful invasions by new species and the growth rate of coexisting cell species. (B) Plot of leakage (blue) and uptake (red) fluxes of non-nutrient chemicals from each cell species A-F. (C) Structure of metabolic exchange among six cell species that have different growth rates in isolation. The vertical axis represents the growth rate of each cell species ***α*** in isolation, 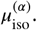 Cyan and pink arrows indicate the leakage and uptake of each chemical component, respectively. Symbiosis among multiple species increases the growth rate to ***μ***_symbiosis_ (as indicated on the top), which is higher than the growth rate of each cell species in isolation, 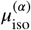. In the numerical simulation, the parameters were set to ***n*** = 20, ***S***_env_ = 0.03, ***V***_env_ = 3, 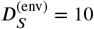, *D_S_* = 1, ***R***_deg_ = 5 × 10^-5^, ***N***_enzyme_ = ***n***/5.

Fig. 3BC also shows that every cell species leaks some metabolites and consumes others, and metabolites are exchanged between all cells as a result of cell-level adaptation. Unlike the assumptions of food chains or BQH (***Morris et al., 2012**; **Morris, 2015**; **Goyal and Maslov, 2018***), the leakerconsumer relations via metabolic exchanges are entangled, and not hierarchical or cyclic. Hence, no clear trophic levels are observed.

### Cell-level adaptation via leak advantage frequently leads to symbiosis

Wethen statistically examined how frequently such symbiosis was realized (Fig. 4). Figure 4A shows the frequency of symbiotic coexistence of multiple species in a single-nutrient condition. As the number of chemical components ***n*** increases, the coexistence of more species is more likely. For ***n*** = 30, symbiotic coexistence is achieved for almost all the trials, as long as the cell species with the fastest growth in isolation has at least one leak-advantage chemical. Here, recall that the frequency of leak advantage in isolation increases with ***n*** (***Yamagishi et al., 2020***).

**Figure 4.**
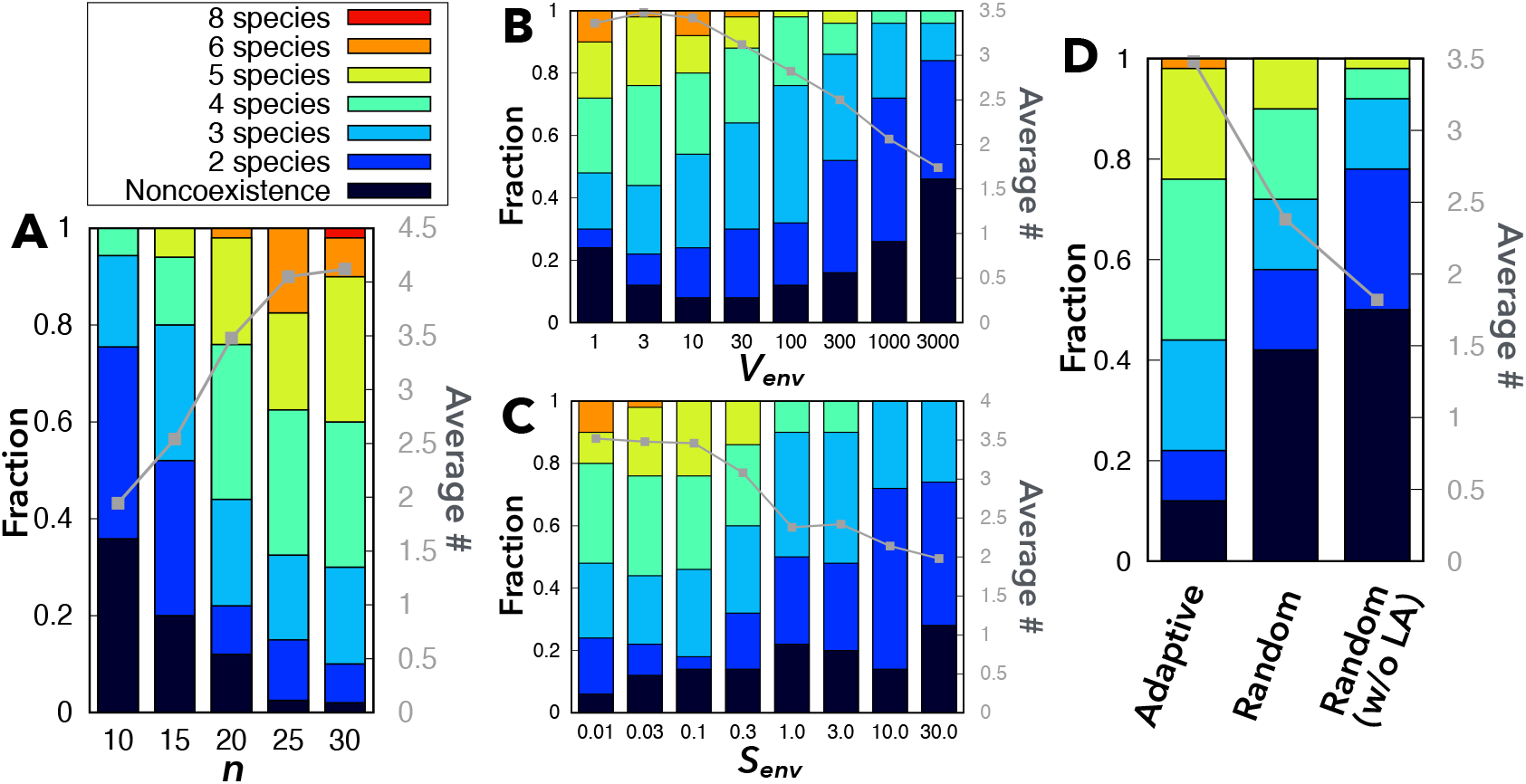
Statistics of symbiosis among randomly generated networks. 50 independent trials were conducted for each set of parameters. (A) Dependence of the frequency of coexisting species on the number of chemical components *n*. ***S***_env_ = 0.03, ***V***_env_ = 3. (B) Dependence of the frequency of coexisting species upon ***V***_env_. *n* = 20, ***S***_env_ = 0.03. (C) Dependence of the frequency of coexisting species upon ***S***_env_. *n* = 20, ***V***_env_ = 3. (D) The frequency of coexisting species for random fixed diffusion coefficients (with and without leakage of chemicals that confer leak advantage) without cell-level adaptation. *n* = 20, ***S***_env_ = 0.03, ***V***_env_ = 3. In the “random” case, the diffusion coefficients of chemicals ***N***_enzyme_ +1 ~ *n* – 1 are chosen randomly from a uniform distribution [0.0: 1.0] (see also Fig. S6). In all the figures, the colored bars show the frequency of symbiosis among two to eight species (with different colors), whereas the black bars show noncoexistence. The frequency for each parameter set was calculated from 50 independent samples of *N* catalytic networks where the species with the fastest growth in isolation has a leak-advantage chemical in its reaction network. In all the numerical simulations, the other parameters are fixed: 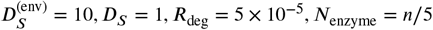.

In addition, symbiosis is achieved frequently with a wide range of environmental parameters ***S***_env_ and ***V***_env_ (and ***R***_deg_) (Fig. 4BC and Fig. S4). Notably, Fig. 4B demonstrates that the frequency of symbiosis decreases as the size of the environment ***V***_env_ increases, unless the environment is too small (***V***_env_ ≃ 1, i.e., the total volume of cells equals that of the environment). Note here that increase in ***V***_env_ (and/or ***R***_deg_) weakens cell-cell interactions because the secreted chemicals are diluted; consequently, the growth change due to consumption is suppressed while that due to leakage is little affected. Hence, 1/***V***_env_ (and 1/***R***_deg_) serves as an indicator of the strength of cell-cell interactions (or the efficiency of exchange of secreted metabolites). Indeed, for larger ***V***_env_, symbiosis is achieved less frequently by metabolite exchange via the environment. Note that when ***V***_env_ is too small (***V***_env_ ≃ 1), the environmental concentration of chemicals is sensitive to addition of new species, and therefore, the coexistence of multiple species becomes unstable.

Figure 4C shows that decrease in the nutrient supply ***S***_env_ results in increase in the frequency of symbiosis and the average number of coexisting species. This may be explained by the increasing importance of metabolic efficiency of converting the nutrient to biomass on growth rate in large ***S***_env_, relative to the efficiency of the exchange of non-nutrient metabolites. Hence, cell species with slower growth in isolation (which is based on metabolic efficiency of the nutrient) may not be able to achieve the same growth rate as those with faster growth in isolation.

In short, the results shown in Fig. 4A-C suggest that symbiosis among multiple species via adaptive metabolite exchange is commonly achievable because microbes contain many chemical components and often exist in crowded and nutrient-poor environments.

### Decrease in species diversity in the absence of cell-level adaptation

Furthermore, to ascertain the contribution of cell-level adaptation to symbiosis (via leak advantage), we also considered situations where the leakage and uptake of metabolites occur simply because of the inevitable permeability of cellular membranes, and compared them with the results under the assumption of cell-level adaptation. In this case, the diffusion coefficients of invading cells are given fixed, random positive values for all metabolites (Fig. S5). Interestingly, the frequency of coexistence of multiple species is much smaller (Fig. 4D).

Notably, when adaptive changes in the diffusion coefficients at the cell level are allowed, leakage occurs only when it promotes the growth of the leaker cell species (otherwise, the leaker cells would decrease their diffusion coefficient to zero). A mutualistic relationship is thus necessarily established between leaker and consumer cells. Mutualistic cell-cell interaction usually leads to stable coexistence, as discussed in previous studies (***Tokita and Yasutomi, 2003**; **Pande et al., 2014**; **Giri et al., 2020***). In contrast, leakage of the metabolites by random fixed diffusion coefficients does not necessarily benefit leaker cells, thereby making leaker-consumer interactions often parasitic or even competitive (see Fig. S6B). Such parasitic or competitive interactions often make the steady state (linearly) unstable (***May, 1971**; **Hassell and May, 1973***). Consistently, for the case with randomly pre-fixed diffusion coefficients, the frequency of parasitic (symbiotic) relationships decreases (increases) as the number of coexisting species increases(Fig. S6B). Hence, the likelihood of coexistence is lower when the diffusion coefficients are fixed.

To further corroborate these results, we also generated a model with null diffusion coefficients for chemicals that confer leak advantage (in isolated conditions) and positive random coefficients for chemicals that do not. In this situation, leakage is always disadvantageous in isolated conditions. In the absence of leakage of leak-advantage chemicals, the frequency of symbiosis and the average number of coexisting species were further reduced (Fig. 4D), even though growth promotion due to the uptake of metabolites could still occur.

### Resilience of symbiosis mediated by metabolite exchange

Thus far, we investigated if and how symbiosis of diverse cell species with tangled forms of metabolite exchange is achieved as a result of cell-level adaptation of diffusion coefficients. Lastly, we examined the stability of communities consisting of diverse cell species that exchange metabolites.

In particular, we focused on the resilience of symbiotic relationships against the removal of a species that coexisted. In most cases, the removal of one species from the community does not cause the successive extinction of any other species in the community (Fig. 5A). In contrast to the extinction of species in a hierarchical ecosystem with trophic levels, where the removal of some *keystone/core species* leads to an *avalanche* of extinctions of downstream species in the hierarchy (***Paine, 1969; Mills et al., 1993; Goyal and Maslov, 2018***), such an avalanche of species extinctions hardly occurs in our model (Fig. 5A). In other words, we rarely observe the existence of keystone/core species whose absence prevents many other species from coexisting, and removal of which leads to the extinction of all other species. Hence, the present system with metabolite exchange has a high degree of resilience.

**Figure 5.**
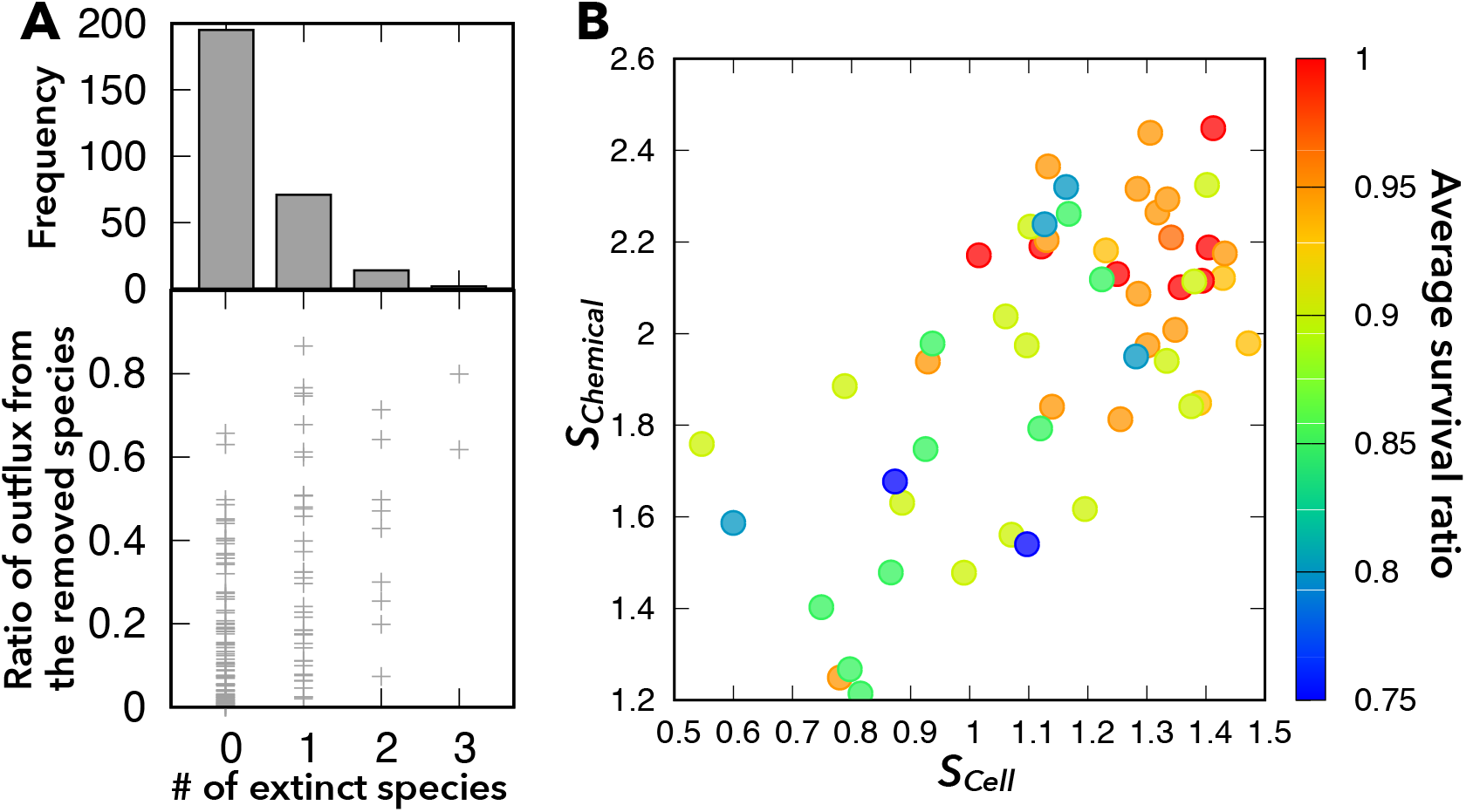
Resilience of symbiotic coexistence against the removal of one species. (A) The upper panel shows the frequency distribution of the number of cell species (0-3) that become extinct when one species is removed. The lower panel shows the ratio of leakage of chemicals from the removed cell species to the total leakage from all cell species to the environment. Each point corresponds to samples shown in the upper panel. (B) Average survival ratio (color) against the indices that characterize the diversity of leaking cell species (***S***_Cell_) and leaked chemical components (***S***_Chem_). The survival ratio is the number of surviving species after removal of one species, divided by the number of species before the extinction successive to the removal. (If no additional cell species become extinct, this ratio equals one.) The multiple correlation coefficient between survival ratio and (***S***_Cell_, ***S***_Chem_) is 0.49, while the correlation coefficients between the survival ratio and ***S***_Cell_, between the survival ratio and ***S***_Chem_, and between ***S***_Cell_ and ***S***_Chem_ are 0.46, 0.43, and 0.65, respectively. For these calculations, we used 55 samples of coexistence of five or six species with ***n*** = 20.

As shown in Fig. 5A, we did observe a few cases in which removal of one species causes the extinction of most species. In such non-resilient cases, the removed species tended to be ones that dominantly leaked chemicals into the environment. In general, the resilience of the system to removal of species increased with the extent of entanglement of metabolite exchange across cells (i.e., many cell species leak many chemical components) (see also Fig. S7). To quantitatively characterize this tendency, we introduced the indices of the effective “entropies” that characterize the diversity of leaking cell species and leaked chemical components as

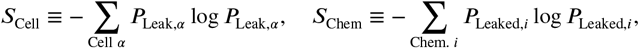

where the total flux of all chemicals from cell species ***α*** and the total flux of chemical ***i*** from all cell species to the environment are defined as 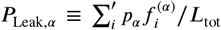 and 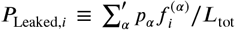, respectively, with 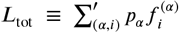, where the summation ∑’ is taken only for leakage (not for uptake), that is 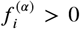. If ***N*** cell species leak chemicals evenly, ***S***_Cell_ takes log*N* (maximal value), and if ***n*** chemicals are leaked evenly, ***S***_Chem_ takes log ***n***. In contrast, if only one cell species leaks or only one chemical is leaked, ***S***_cell_ or ***S***_Chem_ is equal to 0.

Indeed, when a large number of cells leak and exchange a large number of chemical components (i.e., both ***S***_Cell_ and *S*_Chem_ are large), the system tends to be resilient (Fig. 5B). ***S***_Cell_ and *S*_Chem_ are also strongly correlated (Fig. 5B), such that when many cell species contribute to leakage, many components are leaked, and vice versa. We now consider two extreme situations to illustrate this correlation: if ***S***_Chem_ = 0 (i.e., only one chemical is leaked to the environment), then *S*_Cell_ must be 0 due to Gause’s rule. In contrast, when ***S***_Chem_ is large, ***S***_Cell_ is unlikely to be zero because a large ***S***_Chem_ allows the coexistence of many cells, which in turn can leak more chemicals, leading to large ***S***_Cell_ in steady states.

## Discussion

In this paper, we elucidated how symbiosis mediated by the exchange of various metabolites among diverse cell species is possible, based on the advantages of metabolite leakage for leaker cells. Resilient symbiosis among diverse species can be achieved when each cell species is allowed to adaptively change the degree of leakage and uptake of metabolites.

First, we described the mechanism and conditions for mutualism between leaker and consumer cells. As the density of cells (with leak advantage) is increased, the metabolites secreted by the leaker cells accumulate in the environment, thereby preventing further leakage. Consequently, coexistence with a different species that consumes the leaked metabolite for its own growth brings about further leak advantage for the leaker cells, while the consumption of the leaked metabolite is beneficial for the growth of the consumer cells. In this leaker-consumer mutualism, both cells increase their growth rates through cell-cell interactions mediated by secreted metabolites.

In contrast, whether the leakage is beneficial for the leaker cells is not fully addressed in the previous studies on microbial ecology (***Morris et al., 2012; Morris, 2015; Großkopf et al., 2016; Zomorrodi and Segrè, 2017***). Though the BQH also discusses the evolution of cooperation (***Morris, 2015; Sachs and Hollowell, 2012***), it often (implicitly) assumes that metabolic secretion leads to parasitism or free riding, as the leakage is just an inevitable consequence of a permeable membrane and thus the leakage is not necessarily advantageous for the leaker cells. Although this assumption is not unreasonable and is consistent with some empirical observations (***Morris, 2015; Gore et al., 2009; Wang and Goldenfeld, 2011***), it is not clear why the leaker cells have not evolved mechanisms to suppress such leakage that may be disadvantageous. In this respect, our results will complement the BQH: some microbial cells secrete chemicals just because this process is beneficial for them. In this sense, the “richer” cells “donate” their products to “poorer” cells for the sake of the former cells themselves, as if the cells are practicing “potlatch” often seen in human societies (***Mauss, 1970**; **Bataille and Hurley, 1988***).

This “microbial potlatch” generally emerges as a result of individual-level adaptation under conditions of complex intracellular metabolic networks, crowded environments, and limited nutrient supply. Indeed, facilitation of growth due to the coexistence of different strains or species has been reported in several experiments (***Ponomarova et al., 2017; Wintermute and Silver, 2010; Kosina et al., 2016; Pande et al., 2014***) and in laboratory evolution under nutrient depletion (***Hillesland and Stahl, 2010***). Although some studies have reported that lower resource availability leads to a more diverse community (***Zhou et al., 2002; Rajaniemi, 2003***), other studies have reported contradictory results (***Waldrop et al., 2006; Øvreås and Torsvik, 1998***). Of note, the existence of waste byproducts and inhibitory chemicals, which was not assumed, could lead to greater diversity under nutrient-rich environments, as greater nutrient supply will increase the waste byproducts (***Pfeiffer and Bonhoeffer, 2004; Marsland III et al., 2019***). Indeed, the actual relationship between diversity and resource availability is sometimes non-monotonous (***Claire Horner-Devine et al., 2003; Mancuso et al., 2020***) and can depend on the cultivation conditions (***Zhou et al., 2002***). This complex relationship may be a consequence of the competing effects of essential metabolites and waste byproducts.

Moreover, the symbiotic coexistence of species is seen more often when diffusion coefficients are not fixed and cell-level adaptation is allowed. When diffusion coefficients are fixed, even though all metabolites are leaked into the environment and thus the number of niches is larger, we found that the actual number of coexisting species in our simulations is much lower than that in the case with adaptive changes in diffusion coefficients (and is also lower than the Gause’s limit). Leaker-consumer mutualism by cell-level adaptation spontaneously reaches optimal values of diffusion coefficients that facilitate high growth rates of all cell species and allow for stable coexistence.

Furthermore, we examined the ecological resilience of microbial communities against the removal of a coexisting species. As more cells leak and exchange more chemical components in an ecosystem (i.e., ***S***_Cell_ and ***S***_Chem_ are large), the microbial community becomes more resilient to the removal of its members. In contrast, if only few cell species leak a few chemicals, the ecosystem would have a unidirectional structure similar to a “food chain.” In such cases, keystone/core species could exist, and the system would not be resilient to removal of such species. Empirical studies suggest that diverse microbial communities are more resistant to environmental disturbances than monocultures (***Giri et al., 2020***). Experimentally estimating ***S***_Cell_ or ***S***_Chem_ in these microbial communities and investigating their relationship with their resilience could be interesting.

Theoretically, we find that coexistence of diverse species is achieved by incorporating multilevel dynamics at the intercellular (population) and intracellular (metabolic) levels and cannot be captured by standard Lotka-Volterra-type population dynamics. Microbial ecosystems with metabolite exchange via the environment are expected to behave differently from those with simple food chain or food web structures that are often considered in Lotka-Volterra-type population dynamics, as the interactions between different cell species depend not only on their populations but also on the exchanged chemicals, which depend on their intracellular states (***Liao et al., 2020***). For example, in leaker-consumer mutualism, the benefit for leaker cells is indirect; the leaker cells benefit from the consumption of accumulated chemicals only when the density of leaker cells is high enough to cause excess accumulation of these chemicals. The leaker-consumer mutualism is thus frequency-dependent and depends on the degree of interaction between cells via secreted chemicals. In the present model, this degree of cell-cell interaction depends on the relative volume of the medium to that of a cell ***V***_env_ (i.e., the inverse of cell density) and the degradation rate of chemicals in the medium ***R***_deg_. If ***V***_env_ and ***R***_deg_ are sufficiently large, the leaker cells can continue to leak chemicals and grow efficiently without consumer cells. In this sense, leaker-consumer mutualism with unidirectional flow is different from ordinary forms of cooperation or division of labor (***Yamagishi et al., 2016; Flores and Herrero, 2010***). Still, in a system with multiple cell species we studied here, each cell can simultaneously be a leaker and a consumer for different chemicals, thus achieving metabolic division of labor with bidirectional exchange.

The premise of the present study was that the leakage of even essential metabolites can be beneficial for cellular growth under certain conditions, and the control of leakage provides a possible means of adaptation. In principle, this theory of leak advantage is testable through experiments on microbes where the concentration of some secreted chemicals in the culture medium is fixed by means of a chemostat and the dependence of the cellular growth rate on the extracellular concentration is measured.

Indeed, the coexistence of multiple species via active secretion of chemicals is considered in the context of classical syntrophy in microbial communities (***Morris et al., 2013**; **Cavaliere et al., 2017***), where it is generally assumed that the leaked chemicals are non-essential or inhibitory to the leaker itself, but are useful for the other species. Such chemicals could surely exist but the leaked essential chemicals would be likely to be useful for more diverse species (which would allow for the coexistence of more diverse species). Moreover, the leaked chemicals in classical syntrophy are thought to be located ata lower level of a chemical hierarchy according to energetics (***Embree et al., 2015; Morris et al., 2013***), which would also determine a trophic hierarchy. In contrast, chemicals that confer leak advantage are often essential, and lead to entangled networks of metabolite exchanges between different cell species, as often observed in actual microbial ecosystems (***Goldford et al., 2018; Baran et al., 2015; Zengler and Zaramela, 2018***).

Finally, let us discuss whether a leak advantage would be eliminated through the course of evolution by incorporating appropriate gene-regulation of enzymatic activity, which is a well-known means to optimize cell growth (***Jacob and Monod, 1961***). If only a single cell species exists, such optimization of gene regulation might be possible to eliminate the leak advantage by evolution. However, if cells interact with other cells, it would be difficult to reach such optimized growth state without leakiness, through evolutionary optimization in enzymatic activity. As the cell numbers increase, the environment inevitably becomes crowded, and cell-cell interactions through secreted chemicals cannot be disregarded. Hence, optimization cannot occur under isolated conditions unless cells find an optimized solution without any secretion of chemicals. Once the evolution progresses under the environmental conditions with interacting cells, finding such a solution, even if it exists, would take many generations. Before such isolated optimization is reached, other cells that consume secreted chemicals could either emerge through mutation or invade from elsewhere, thereby enhancing the growth of the leaker species. Then, symbiotic relationships with different cell species will develop and result in further entanglement of chemical exchange networks, as described in this paper. Indeed, some experiments have shown the *de novo* emergence of coexistence via metabolite secretion (***Ponomarova et al., 2017; Le Gac et al., 2012; Hom and Murray, 2014***), and nonspecific metabolic cross feeding has been reported to lead to the coexistence of different phenotypes in such a community (***Goldford et al., 2018; Ponomarova and Patil, 2015***). Moreover, a recent study on phylogeny suggested that such leakage of essential metabolites has been promoted through evolution and adaptation (***Braakman et al., 2017***).

In summary, we have shown that cell-level adaptation of leakiness of (essential) metabolites spontaneously establishes symbiotic relationships. This “microbial potlatch” generally emerges when the intracellular metabolic network is complex, the environment is crowded, and nutrient supply is limited. The present study provides a basis for complex microbial ecosystems with diverse species.

## Acknowledgments

The authors would like to thank Chikara Furusawa and Kazufumi Hosoda for their useful comments. This research was partially supported by a Grant-in-Aid for Scientific Research (A) (20H00123) and Grant-in-Aid for Scientific Research on Innovative Areas (17H06386) from the Ministry of Education, Culture, Sports, Science and Technology (MEXT) of Japan.

## References

Baran R, Brodie EL, Mayberry-Lewis J, Hummel E, Da Rocha UN, Chakraborty R, Bowen BP, Karaoz U, Cadillo-Quiroz H, Garcia-Pichel F, et al. Exometabolite niche partitioning among sympatric soil bacteria. Nature communications. 2015; 6:82–89.

Bataille G, Hurley R. The accursed share, vol. 1. Zone Books New York; 1988.

Braakman R, Follows MJ, Chisholm SW. Metabolic evolution and the self-organization of ecosystems. Proceedings of the National Academy of Sciences. 2017; 114(15):E3091–E3100.

Cavaliere M, Feng S, Soyer OS, Jiménez JI. Cooperation in microbial communities and their biotechnological applications. Environmental microbiology. 2017; 19(8):2949–2963.

Claire Horner-Devine M, Leibold MA, Smith VH, Bohannan BJ. Bacterial diversity patterns along a gradient of primary productivity. Ecology letters. 2003; 6(7):613–622.

Curtis TP, Sloan WT, Scannell JW. Estimating prokaryotic diversity and its limits. Proceedings of the National Academy of Sciences. 2002; 99(16):10494–10499.

Datta MS, Sliwerska E, Gore J, Polz MF, Cordero OX. Microbial interactions lead to rapid micro-scale successions on model marine particles. Nature communications. 2016; 7(1):1–7.

D’Souza G, Waschina S, Pande S, Bohl K, Kaleta C, Kost C. Less is more: selective advantages can explain the prevalent loss of biosynthetic genes in bacteria. Evolution. 2014; 68(9):2559–2570.

Embree M, Liu JK, Al-Bassam MM, Zengler K. Networks of energetic and metabolic interactions define dynamics in microbial communities. Proceedings of the National Academy of Sciences. 2015; 112(50):15450–15455.

Flores E, Herrero A. Compartmentalized function through cell differentiation in filamentous cyanobacteria. Nature Reviews Microbiology. 2010; 8(1):39–50.

Furusawa C, Kaneko K. Emergence of rules in cell society: differentiation, hierarchy, and stability. Bulletin of Mathematical Biology. 1998; 60(4):659–687.

Gause GF. Experimental studies on the struggle for existence: I. Mixed population of two species of yeast. Journal of experimental biology. 1932; 9(4):389–402.

Giri S, Shitut S, Kost C. Harnessing ecological and evolutionary principles to guide the design of microbial production consortia. Current Opinion in Biotechnology. 2020; 62:228–238.

Goldford JE, Lu N, Bajić D, Estrela S, Tikhonov M, Sanchez-Gorostiaga A, Segrè D, Mehta P, Sanchez A. Emergent simplicity in microbial community assembly. Science. 2018; 361(6401):469–474.

Gore J, Youk H, Van Oudenaarden A. Snowdrift game dynamics and facultative cheating in yeast. Nature. 2009; 459(7244):253–256.

Goyal A, Maslov S. Diversity, stability, and reproducibility in stochastically assembled microbial ecosystems. Physical Review Letters. 2018; 120(15):158102.

Großkopf T, Consuegra J, Gaffé J, Willison JC, Lenski RE, Soyer OS, Schneider D. Metabolic modelling in a dynamic evolutionary framework predicts adaptive diversification of bacteria in a long-term evolution experiment. BMC evolutionary biology. 2016; 16(1):1–15.

Hardin G. The competitive exclusion principle. Science. 1960; 131(3409):1292–1297.

Hassell MP, May RM. Stability in insect host-parasite models. TheJournal of Animal Ecology. 1973; p. 693–726.

Hillesland KL, Stahl DA. Rapid evolution of stability and productivity at the origin of a microbial mutualism. Proceedings of the National Academy of Sciences. 2010; 107(5):2124–2129.

Hom EF, Murray AW. Niche engineering demonstrates a latent capacity for fungal-algal mutualism. Science. 2014; 345(6192):94–98.

Huber H, Küper U, Daxer S, Rachel R. The unusual cell biology of the hyperthermophilic Crenarchaeon Ignicoc-cus hospitalis. Antonie Van Leeuwenhoek. 2012; 102(2):203–219.

Jacob F, Monod J. Genetic regulatory mechanisms in the synthesis of proteins. Journal of molecular biology. 1961; 3(3):318–356.

Kaneko K. A Scenario for the Origin of Multicellular Organisms: Perspective from Multilevel Consistency Dynamics. In: Niklas KJ, Newman SA, editors. Multicellularity: Origins and Evolution Cambridge: MIT Press; 2016. p. 201–224.

Kaneko K, Yomo T. Cell division, differentiation and dynamic clustering. Physica D: Nonlinear Phenomena. 1994; 75(1-3):89–102.

Kashiwagi A, Noumachi W, Katsuno M, Alam MT, Urabe I, Yomo T. Plasticity of fitness and diversification process during an experimental molecular evolution. Journal of molecular evolution. 2001; 52(6):502–509.

Kosina SM, Danielewicz MA, Mohammed M, Ray J, Suh Y, Yilmaz S, Singh AK, Arkin AP, Deutschbauer AM, Northen TR. Exometabolomics assisted design and validation of synthetic obligate mutualism. ACS synthetic biology. 2016; 5(7):569–576.

Le Gac M, Plucain J, Hindré T, Lenski RE, Schneider D. Ecological and evolutionary dynamics of coexisting lineages during a long-term experiment with Escherichia coli. Proceedings of the National Academy of Sciences. 2012; 109(24):9487–9492.

Liao C, Wang T, Maslov S, Xavier JB. Modeling microbial cross-feeding at intermediate scale portrays community dynamics and species coexistence. arXiv preprint arXiv:200208433. 2020;.

Lozupone CA, Stombaugh JI, Gordon JI, Jansson JK, Knight R. Diversity, stability and resilience of the human gut microbiota. Nature. 2012; 489(7415):220–230.

MacArthur R, Levins R. Competition, habitat selection, and character displacement in a patchy environment. Proceedings of the National Academy of Sciences of the United States of America. 1964; 51(6):1207.

Mancuso CP, Lee H, Abreu CI, Gore J, Khalil AS. Environmental fluctuations reshape an unexpected diversitydisturbance relationship in a microbial community. bioRxiv. 2020;.

Marsland III R, Cui W, Goldford J, Sanchez A, Korolev K, Mehta P. Available energy fluxes drive a transition in the diversity, stability, and functional structure of microbial communities. PLoS computational biology. 2019; 15(2):e1006793.

Mauss M. The gift: Form and functions of exchange in archaic societies. Cohen and West; 1970.

May RM. Stability in multispecies community models. Mathematical Biosciences. 1971; 12(1-2):59–79.

Mills LS, Soulé ME, Doak DF. The keystone-species concept in ecology and conservation. BioScience. 1993; 43(4):219–224.

Morris BE, Henneberger R, Huber H, Moissl-Eichinger C. Microbial syntrophy: interaction for the common good. FEMS microbiology reviews. 2013; 37(3):384–406.

Morris JJ. Black Queen evolution: the role of leakiness in structuring microbial communities. Trends in Genetics. 2015;31(8):475–482.

Morris JJ, Lenski RE, Zinser ER. The Black Queen Hypothesis: evolution of dependencies through adaptive gene loss. MBio. 2012; 3(2).

Øvreås L, Torsvik V. Microbial diversity and community structure in two different agricultural soil communities. Microbial ecology. 1998; 36(3-4):303–315.

Paczia N, Nilgen A, Lehmann T, Gätgens J, Wiechert W, Noack S. Extensive exometabolome analysis reveals extended overflow metabolism in various microorganisms. Microbial cell factories. 2012; 11(1):122.

Paine RT. A note on trophic complexity and community stability. The American Naturalist. 1969; 103(929):91–93.

Pande S, Merker H, Bohl K, Reichelt M, Schuster S, De Figueiredo LF, Kaleta C, Kost C. Fitness and stability of obligate cross-feeding interactions that emerge upon gene loss in bacteria. The ISME journal. 2014; 8(5):953–962.

Pfeiffer T, Bonhoeffer S. Evolution of cross-feeding in microbial populations. The American naturalist. 2004; 163(6):E126–E135.

Ponomarova O, Gabrielli N, Sévin DC, Mülleder M, Zirngibl K, Bulyha K, Andrejev S, Kafkia E, Typas A, Sauer U, et al. Yeast creates a niche for symbiotic lactic acid bacteria through nitrogen overflow. Cell systems. 2017; 5(4):345–357.

Ponomarova O, Patil KR. Metabolic interactions in microbial communities: untangling the Gordian knot. Current opinion in microbiology. 2015; 27:37–44.

Rajaniemi TK. Explaining productivity-diversity relationships in plants. Oikos. 2003; 101(3):449–457.

Rosenzweig RF, Sharp R, Treves DS, Adams J. Microbial evolution in a simple unstructured environment: genetic differentiation in Escherichia coli. Genetics. 1994; 137(4):903–917.

Sachs J, Hollowell A. The origins of cooperative bacterial communities. MBio. 2012; 3(3).

Tokita K, Yasutomi A. Emergence of a complex and stable network in a model ecosystem with extinction and mutation. Theoretical population biology. 2003; 63(2):131–146.

Waldrop MP, Zak DR, Blackwood CB, Curtis CD, Tilman D. Resource availability controls fungal diversity across a plant diversity gradient. Ecology Letters. 2006; 9(10):1127–1135.

Wang Z, Goldenfeld N. Theory of cooperation in a micro-organismal snowdrift game. Physical Review E. 2011; 84(2):020902.

Wintermute EH, Silver PA. Emergent cooperation in microbial metabolism. Molecular systems biology. 2010; 6(1):407.

Yamagishi JF, Saito N, Kaneko K. Symbiotic cell differentiation and cooperative growth in multicellular aggregates. PLoS computational biology. 2016; 12(10):e1005042.

Yamagishi JF, Saito N, Kaneko K. Advantage of Leakage of Essential Metabolites for Cells. Physical Review Letters. 2020; 124(4):048101.

Zelezniak A, Andrejev S, Ponomarova O, Mende DR, Bork P, Patil KR. Metabolic dependencies drive species co-occurrence in diverse microbial communities. Proceedings of the National Academy of Sciences. 2015; 112(20):6449–6454.

Zengler K, Zaramela LS. The social network of microorganisms—how auxotrophies shape complex communities. Nature Reviews Microbiology. 2018; 16(6):383–390.

Zhou J, Xia B, Treves DS, Wu LY, Marsh TL, O’Neill RV, Palumbo AV, Tiedje JM. Spatial and resource factors influencing high microbial diversity in soil. Applied and Environmental Microbiology. 2002; 68(1):326–334.

Zomorrodi AR, Segrè D. Genome-driven evolutionary game theory helps understand the rise of metabolic interdependencies in microbial communities. Nature communications. 2017; 8(1):1–12.

