## Supplementary Material for "Microbial Potlatch: Cell-level adaptation of leakiness of metabolites leads to resilient symbiosis among diverse cells"

### Details of the Model of Symbiosis among Randomly Generated Networks

To numerically examine how complicated forms of symbiosis can emerge because of cell-level optimization for each cell's growth, the following algorithm is employed (Fig. S1). Here, note that the diffusion coefficient of the nutrient, $D_0^{(\alpha)}$, is fixed to 1.0 for all cell species $\alpha$ throughout the simulation.

**Step 1. Prepare a "species pool" that contains $N = 50$ cell species with different, randomly generated reaction networks.** (see the next subsection for details).

**Step 2. Select the cell species 0 whose growth rate in isolation, $\mu_{\rm iso}^{(0)}$, is highest among the $N$ species.** First, only cell species 0 with the fastest growth in isolation is placed in an environment with fixed $V_{\rm env}$, $S_{\rm env}$, and $D_S^{(\rm env)} = 10$. Then, the diffusion coefficient of each non-nutrient metabolite $i$ is chosen from $D_i^{(0)} = 0.001, 0.002, 0.004, \cdots, 1.024$ to optimize the growth rate of the cell species. If the optimal diffusion coefficients of all non-nutrient metabolites, $D_i^{(0)}$, equal zero (i.e., if species 0 has no leak-advantage chemicals), the species pool is re-generated (otherwise, species 0 dominates and coexistence of multiple specie does not occur).

**Step 3. Cell-level adaptation of the diffusion coefficient for the invading species.** Each cell species with the other $N-1$ networks is examined, one by one, to determine whether they can invade (i.e., grow and coexist) or not. This new cell species $\beta$ initially constitutes a proportion $p_\beta = 0.0025$ of the population, and the diffusion coefficient of each non-nutrient metabolite $i$ is selected from $D_i^{(\beta)} = 0.001, 0.002, 0.004, \cdots, 1.024$ so that the growth rate is maximized in the environmental condition with pre-existing cell species (initially only the fastest growing species, given at Step 2, exists). (In other words, at the onset of the invasion, the invading species' population is small enough not to affect the environmental conditions.) Next, the population dynamics of $\dot{p}_\alpha = (\mu_\alpha - \bar{\mu})p_\alpha$ are computed for a period $T = 1.8 \times 10^6$ that is long enough for populations to reach a stationary distribution with fixed $\mathbf{D}^{(\alpha)}$. Here, if the population ratio $p_\alpha$ of a cell species $\alpha$ is below $p_{\min} \equiv 0.00025$, the cell species $\alpha$ becomes extinct.

**Step 4. Cell-level adaptation of the diffusion coefficient for all coexisting species after invasion.** Each of the surviving species independently changes its diffusion coefficient by $\Delta D_{\text{Leak}} = 0.001$ to optimize its growth over a sufficiently long period $T$ (see the following subsection and Fig. S2 for details). After that, the population dynamics are calculated over a sufficiently long period $T$ without changing their diffusion coefficients $\mathbf{D}^{(\alpha)}$.

**Step 5. Repeat steps 3 and 4 (i.e., invasion and adaptation) until the population of coexisting species converges.**

In this model, it is assumed that the timescales of the following three procedures are distinct: "the dynamical change of the environmental concentration and population" $\ll$ "the adaptation of the diffusion coefficients of each cell species" $\ll$ "the invasion of new species." Assuming this timescale separation, the above algorithm is justifiable as a model of the invasion of the microbial ecological system by additional species.

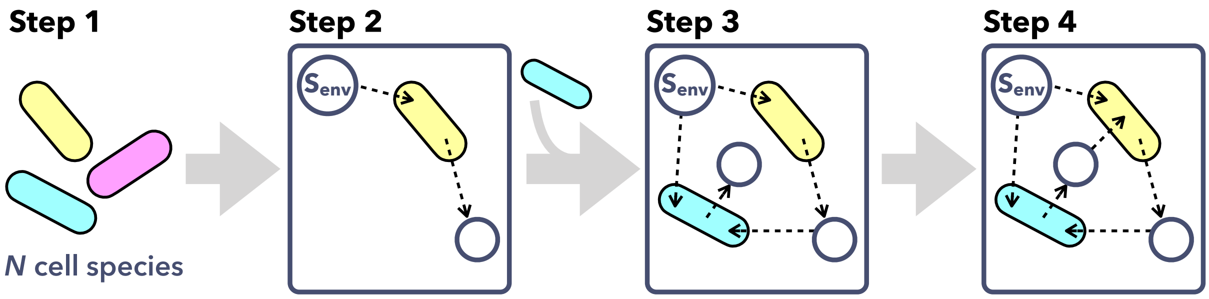


Figure S1: Schematic illustration of the model of symbiosis among randomly generated networks.

### Preparing a "species pool" of randomly generated networks in Step 1

We randomly generated catalytic reaction networks with $n$ kinds of chemical components as shown below (see also Yamagishi *et al.*, 2020). Subsequently, we selected those networks that provide stable growth, namely, those whose growth rate $\mu$ is higher than $5 \times 10^{-5}$.

Each catalytic network consists of $n$ chemicals and $\rho n$ catalytic reactions with $\rho = 2$. Chemicals 1 to $N_{\text{enzyme}}$ are defined as enzymes and they can be a catalyst or product of catalytic reactions but cannot be a substrate of reactions, while the rest of the chemicals (chemicals $N_{\text{enzyme}} + 1$ to $n - 1$ and nutrient chemical 0) are defined as metabolites that can be substrates or products but cannot catalyze any reactions.

Here, the cellular growth rate is determined by the synthesis rate of a single chemical that contributes to biomass, where metabolite $n - 1$ and enzyme 1 are defined as the precursor for biomass synthesis and enzyme for biomass synthesis, respectively. The cellular growth rate $\mu(\mathbf{x}; \mathbf{D}, \mathbf{x}^{(\text{env})})$ is thus given by the rate of biomass synthesis, $\mu(\mathbf{x}) = k_{\text{BM}} x_{n-1} x_1$.

Then, the time course of the concentration of chemical $i$ ($i = 1, 2, \cdots, n-2$) is expressed as

$$\frac{d}{dt} x_i = \underbrace{\sum_{j,l=0}^{n-1} P(j,i,l) x_j x_l - \sum_{j,l=0}^{n-1} P(i,j,l) x_i x_l \quad \underbrace{-D_i x_i}_{f_i(\mathbf{x}; D_i, x_i^{(\text{env})})}}_{F_i(\mathbf{x})} \quad -\mu(\mathbf{x}; \mathbf{D}, \mathbf{x}^{(\text{env})}) x_i,$$

where $P(i,j,l)$ takes the value of one if reaction $i + l \to j + l$ occurs; Otherwise, it equals 0, and all the reaction rate constants are set to unity. For chemicals 0 (nutrient) and $n - 1$ (biomass precursor), evolution can be written as

$$\frac{d}{dt} x_0 = -\sum_{j,l=0}^{n-1} P(0,j,l) x_0 x_l + D_S (S_{\text{env}} - x_0) - \mu(\mathbf{x}; \mathbf{D}, \mathbf{x}^{(\text{env})}) x_0,$$

$$\frac{d}{dt} x_{n-1} = \sum_{j,l=0}^{n-1} P(j, n-1, l) x_j x_l - \sum_{j,l=0}^{n-1} P(n-1, j, l) x_{n-1} x_l - k_{\text{BM}} x_{n-1} x_1 - D_{n-1} x_{n-1} - \mu(\mathbf{x}; \mathbf{D}, \mathbf{x}^{(\text{env})}) x_{n-1}.$$

There are $\rho n$ catalytic reactions, that is, $P(i,j,l)$ has $\rho n$ non-zero elements, which are randomly determined such that a metabolite (i.e., chemical $0, N_{\text{enzyme}} + 1, N_{\text{enzyme}} + 2, \cdots, n - 1$) is transformed to another chemical (an enzyme or a different metabolite, which is not a nutrient 0, i.e., $l \neq 0, i$) by the catalytic action of an enzyme.

### Cell-level adaptation dynamics for diffusion coefficients in Step 4

To numerically compute the cell-level adaptation processes of diffusion coefficients $\mathbf{D}^{(\alpha)}$ (leakage and uptake) by each cell species $\alpha$, we considered the following procedures where each cell species simultaneously increases or decreases the diffusion coefficient of one chemical, say $D_i^{(\alpha)}$, at regular time intervals $\tau$ such that its own growth rate increases.

If the growth rate of cell species $\alpha$ increases after time $\tau$ has passed since $D_i^{(\alpha)}$ is increased (decreased), then $D_i^{(\alpha)}$ is increased (decreased) again; while if the growth rate of cell species $\alpha$ decreases after time $\tau$

has passed since $D_i^{(\alpha)}$ is increased (decreased), then $D_i^{(\alpha)}$ is decreased (increased) (see also Fig. S2):

$$\begin{aligned} D_i^{(\alpha)}(t+\tau+dt) &= D_i^{(\alpha)}(t+\tau) + \Delta D_i^{(\alpha)}(t+\tau) \\ &= D_i^{(\alpha)}(t+\tau) + \Delta D_{\mathrm{Leak}} \mathrm{Sgn}\left[\mu^{(\alpha)}(t+\tau) - \mu^{(\alpha)}(t)\right] \mathrm{Sgn}\left[\Delta D_i^{(\alpha)}(t)\right], \end{aligned}$$

where the initial value is set to $\Delta D_i^{(\alpha)}(t=0) = \Delta D_{\mathrm{Leak}} = 0.001$, and $\tau$ is chosen as $\sim 10/\mu$.

As the diffusion coefficients of all metabolites change adaptively, the index $i$ of the selected chemical is randomly switched to another chemical $i'$ at a constant probability $p_{\mathrm{change}} = 0.1$, if the diffusion coefficient $D_i^{(\alpha)}$ of chemical $i$ has already been changed more than twice. If $D_i^{(\alpha)}(t+\tau) = 0$ and $\Delta D_i^{(\alpha)}(t+\tau) < 0$, then $D_i^{(\alpha)}(t+\tau+dt)$ is maintained at 0, and the index $i$ of the selected chemical is randomly switched to another chemical $i'$.

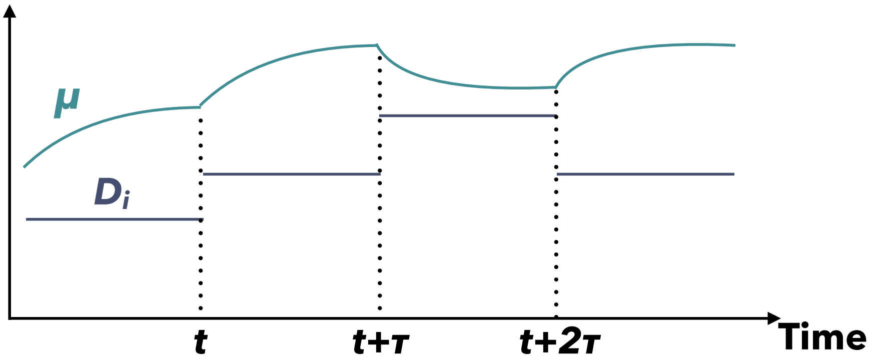


Figure S2: Schematic illustration of cell-level adaptation dynamics of diffusion coefficients. In this example of the time series of diffusion coefficient $D_i^{(\alpha)}$ (dark blue) and $\mu^{(\alpha)}$ (green), $D_i^{(\alpha)}$ is increased at time $t+\tau$ as $D_i^{(\alpha)}(t+\tau+dt) = D_i^{(\alpha)}(t+\tau) + \Delta D_{\mathrm{Leak}}$ because the increase in $D_i^{(\alpha)}$ at time $t$ increases the growth rate as $\mu^{(\alpha)}(t+\tau) > \mu^{(\alpha)}(t)$. Likewise, $D_i^{(\alpha)}$ is decreased at time $t+2\tau$ because the increase in $D_i^{(\alpha)}$ at time $t+\tau$ decreases the growth rate as $\mu^{(\alpha)}(t+2\tau) < \mu^{(\alpha)}(t+\tau)$.

Note that one can consider another way to numerically simulate the cell-level adaptation of the diffusion coefficients. For instance, the growth change due to small additional leakage/uptake can be computed using the inverse matrix of the Jacobi matrix at a steady state [see Eq. [1] in Yamagishi *et al.* (2020) for details], under the assumption that such small changes in diffusion coefficients do not affect the environmental concentrations $\mathbf{x}^{(\mathrm{env})}$. This "more rigorous" procedure for the cell-level adaptation, however, requires much more computing time, while it reproduces qualitatively identical results (Fig. S3).

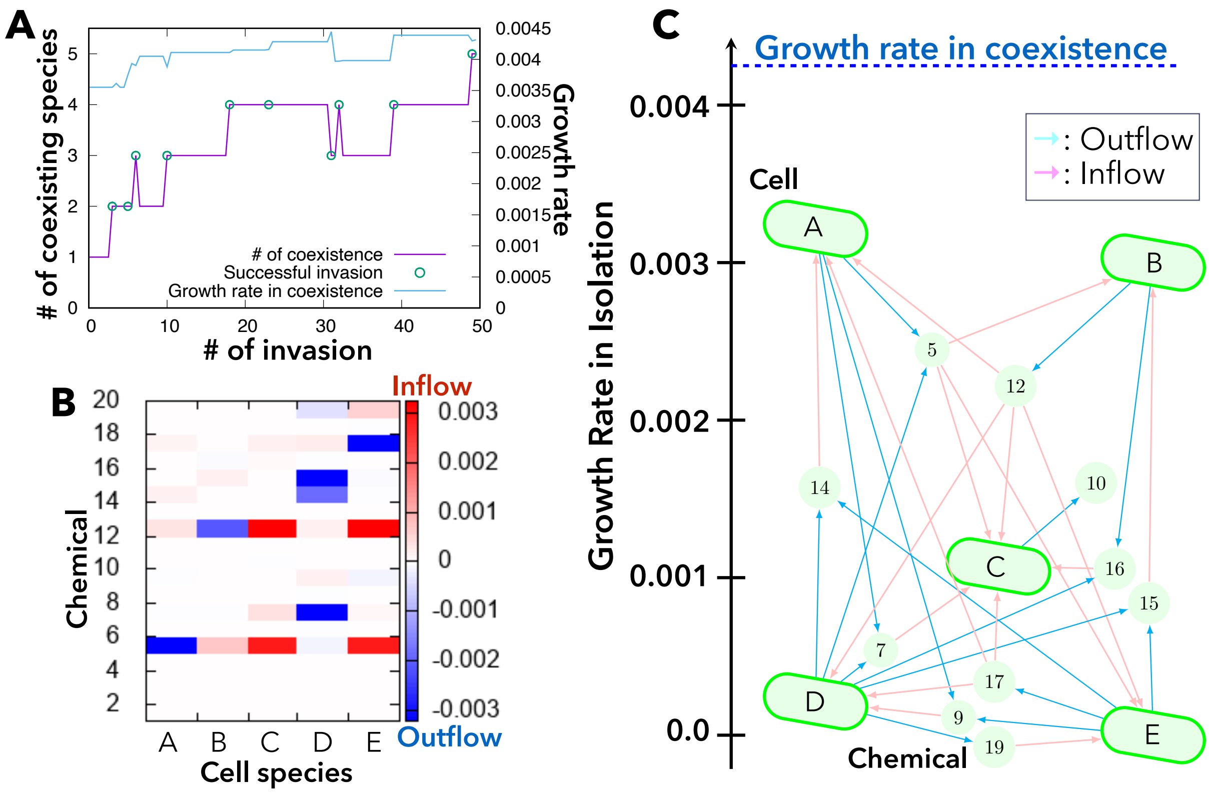


Figure S3: An example of symbiosis when the adaptation dynamics with the Jacobi matrix is adopted. (A) Time series of the number of coexisting species by successful invasions of new species and the growth rate of cell species in coexistence. (B) Plot of leakage (blue) and uptake (red) fluxes of non-nutrient chemicals from each cell species, A-E. (C) Structure of metabolic exchange among five species that originally have different growth rates in isolation. Cyan and pink arrows indicate the leakage and uptake of each chemical component, respectively. Symbiosis among multiple cell species raises the growth rate to reach $\mu_{\text{symbiosis}}$ (as indicated on the top), which is higher than the growth rate of each cell species in isolation, $\mu_{\text{iso}}$. In the numerical simulation, the parameters were set to $n = 20, S_{\text{env}} = 0.03, V_{\text{env}} = 3, D_S^{(\text{env})} = 10, D_S = 1, R_{\text{deg}} = 5 \times 10^{-5}, N_{\text{enzyme}} = n/5$.

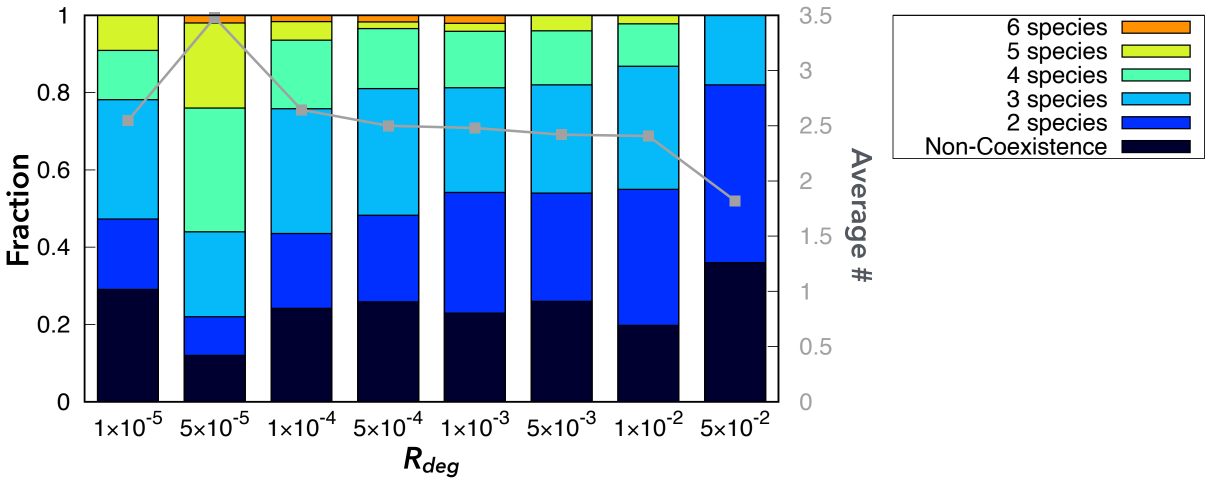


Figure S4: Dependence of the frequency of coexisting species upon $R_{\text{deg}}$. The colored bars show the frequency of symbiosis among two to eight species (with different colors), whereas the black bars show noncoexistence. The frequency for each parameter set was calculated from 50 independent samples of $N$ catalytic networks where the species with the fastest growth in isolation has a leak-advantage chemical in its reaction network. The other parameters are fixed: $n = 20, S_{\text{env}} = 0.03, V_{\text{env}} = 3, D_S^{(\text{env})} = 10, D_S = 1, N_{\text{enzyme}} = n/5$. (see also Fig. 3B in the main text).

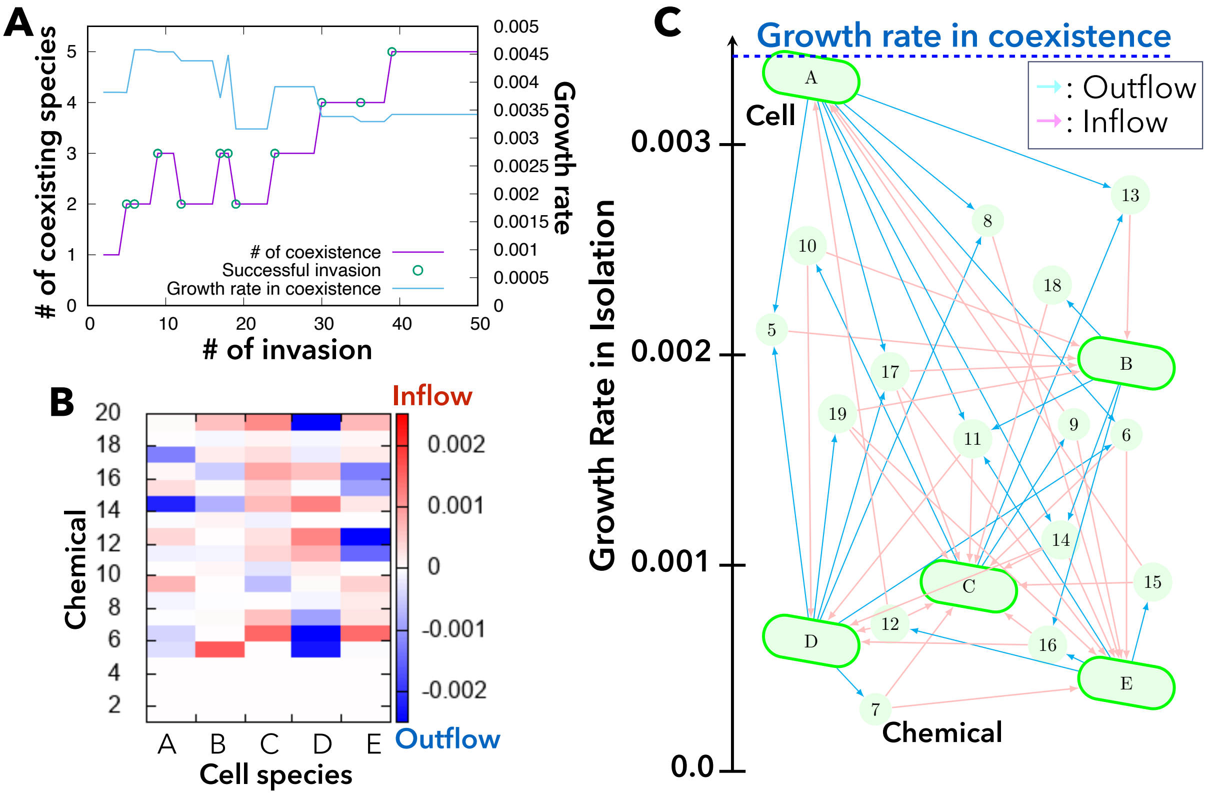


Figure S5: An example of coexistence with randomly pre-fixed diffusion coefficients. The diffusion coefficients of all the non-nutrient metabolites (chemicals $N_{\text{enzyme}} + 1$ to $n - 1$) were randomly chosen from a uniform distribution $[0.0 : 1.0]$. (A) Time series of the number of coexisting species by successful invasions of new species and the growth rate of cell species in coexistence. (B) Plot of leakage (blue) and uptake (red) fluxes of non-nutrient chemicals from each cell species, A-E. (C) Structure of metabolic exchange among five species that originally have different growth rates in isolation. Cyan and pink arrows indicate the leakage and uptake of each chemical component, respectively. The growth rate while coexisting, $\mu_{\text{symbiosis}}$, is indicated on the top blue line. In the numerical simulation, the parameters were set to $n = 20, S_{\text{env}} = 0.03, V_{\text{env}} = 3,\ D_S^{(\text{env})} = 10, D_S = 1, R_{\text{deg}} = 5 \times 10^{-5}, N_{\text{enzyme}} = n/5$.

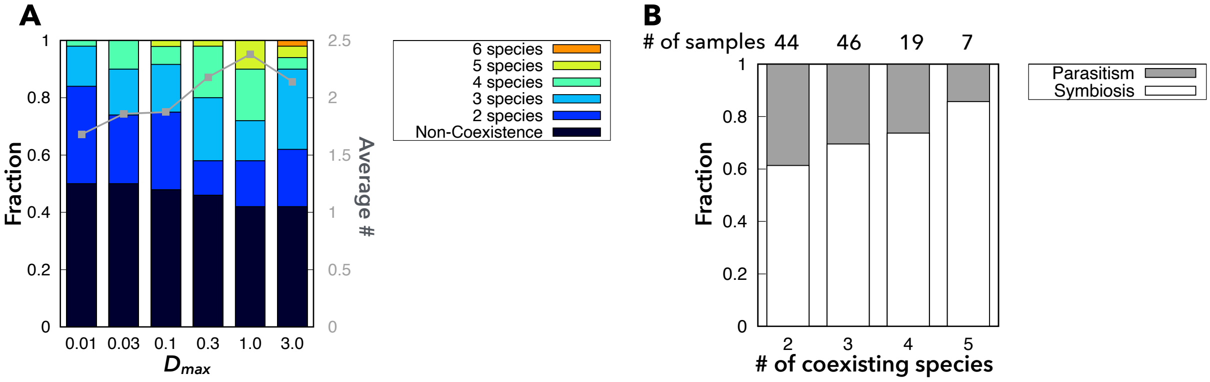


Figure S6: Statistics on coexistence among randomly generated networks with randomly pre-fixed diffusion coefficients. (A) Dependence of the frequency of coexisting species on the upper bound of randomly pre-fixed diffusion coefficients. The diffusion coefficients of non-nutrient metabolites (chemicals $N_{\rm enzyme}+1$ to $n-1$) were randomly chosen from a uniform distribution $[0.0 : D_{\rm max}]$. (B) Relationship between the frequency of symbiosis/parasitism and the number of coexisting species (with various values of $D_{\rm max}$). In all the numerical simulations, the parameters are set at $S_{\rm env} = 0.03, V_{\rm env} = 3, D_S^{(\rm env)} = 10, D_S = 1, R_{\rm deg} = 5 \times 10^{-5}, N_{\rm enzyme} = n/5$.

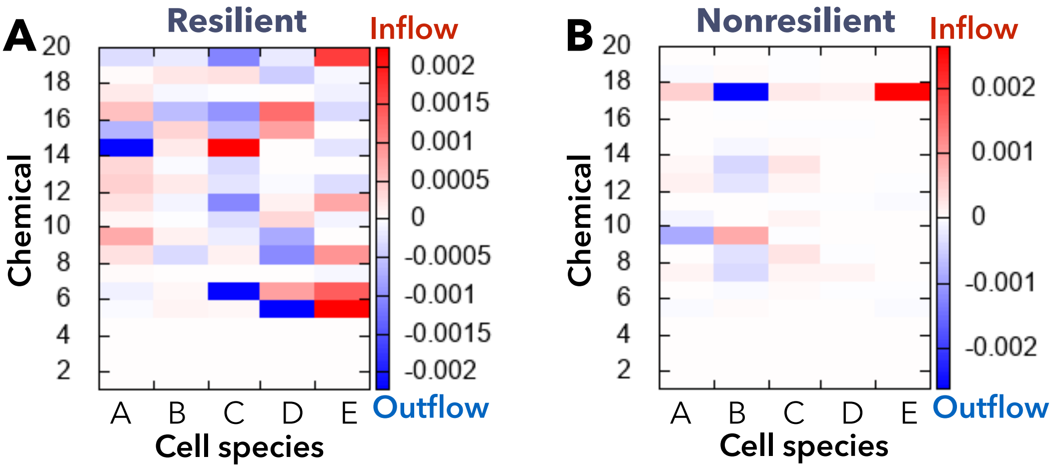


Figure S7: Examples of (A) resilient and (B) non-resilient coexistence. Blue and red indicate the leakage and uptake of each chemical component, respectively. (A) The removal of any species does not cause the extinction of another species (average survival ratio is 1). (B) The removal of cell species B leads to the extinction of the other three cell species (C,D,E) leaving only cell species A in the community (average survival ratio is 0.8). In all the numerical simulations, the parameters are set at $D_S^{(\rm env)} = 10, D_S = 1, R_{\rm deg} = 5 \times 10^{-5}, N_{\rm enzyme} = n/5$.
